# Recent reactivation of a pathogenicity-associated transposable element triggers major chromosomal rearrangements in a fungal wheat pathogen

**DOI:** 10.1101/2023.03.29.534637

**Authors:** Thomas Badet, Alice Feurtey, Daniel Croll

## Abstract

Transposable elements (TEs) are key drivers of genomic variation contributing to recent adaptation in most species. Yet, the evolutionary origins and insertion dynamics within species remain poorly understood. We recapitulate the spread of the pathogenicity-associated *Styx* element across five species that last diverged ∼11,000 years ago. We show that the element likely originated in the *Zymoseptoria* fungal pathogen genus and underwent multiple independent reactivation events. Using a global 900-genome panel of the wheat pathogen *Z. tritici,* we assess *Styx* copy number variation and identify renewed transposition activity in Oceania and South America. We show that the element can mobilize to create additional *Styx* copies in a four-generation pedigree. Importantly, we find that new copies of the element are not affected by genomic defenses revealing a recent loss of control against the element. *Styx* copies are preferentially located in recombination breakpoints and likely triggered multiple types of large chromosomal rearrangements. Taken together, we establish the origin, diversification, and reactivation of a highly active TE with major consequences for chromosomal integrity and the expression of disease.

## Introduction

Transposable elements (TEs) are major constituents of most eukaryotic genomes. Bursts of TE amplification are major drivers of genome evolution as they can create vast adaptive genomic variation (1). However, the repetitive nature of TEs also promotes genome instability through non-allelic homologous recombination potentially triggering chromosomal rearrangements (2). TE mobilization also causes deleterious mutations by the disruption or modification of coding regions that can affect host fitness (3–5). TE sequences relocate to novel loci either directly through DNA or RNA intermediates that often encode the proteins required for sequence excision and integration (1). Highly successful TEs include the short-interspersed nuclear element *Alu*, which is present in >1 million copies in the human genome (6). *Alu*s are non-autonomous elements that hijack the retrotransposition machinery from another retrotransposon for their own mobilization. Similar mechanisms are associated with the recent expansion of the *mPing* miniature-inverted repeat TE in rice (7). Delegating the transposition catalysis to other transcriptionally active elements is an effective strategy for TEs to rapidly increase in copy number. However, retracing the precise movements of TEs across the genome remains challenging due to low sequence complexity of insertion sites and the lack of fully resolved genome assemblies.

A major constraint on TE proliferation in the genome is the activity of the host epigenetic machinery including histone modifications, DNA methylation and RNA interference suppressing active TE copies (8). In rice, both *Ping* and *Pong* can catalyze the transposition of *mPing in vitro,* yet expression of the *Pong* element is effectively repressed by the host silencing machinery so that only the *Ping* element contributes to mobilization (7, 9). To counteract host epigenetic control, some TEs encode their own promoter sequences. The *Ty1*/*Copia*-like retrotransposon *ONSEN* mobilizes upon heat stress in Brassicaceae species and carries a heat-inducible promoter sequence (10, 11). Stress-induced activation has been reported for other TEs across kingdoms and is often associated with transposition bursts (12–15). Activated *ONSEN* elements show an insertion preference for GC- and gene-rich regions (16). Insertion bias towards gene rich sequences likely facilitates the escape from epigenetic control and promotes sustained transposition activity (17).

Independently of transcriptional control, mutations accumulating in TE sequences can dramatically alter transposition rates. Eliminating mutations in the *Sleeping Beauty* Tc1/mariner transposon reactivated the TE in salmon genomes suggesting that these mutations were deleterious to transposon activity (18). In contrast, introducing point mutations at the 3’ terminal inverted repeat (TIR) of the non-autonomous *mPing* element in the rice genome reduces excision events tenfold (19). Ascomycete fungi have evolved a unique mechanism of mutation-driven TE control called repeat-induced point mutations (RIP) (20). Targeted mutagenesis of TE copies disrupts coding sequence integrity through very high rates of C->T transitions. Triggered by the presence of repeated sequences in the genome, RIP can efficiently eliminate the transposition potential of TEs. Understanding how epigenetic factors and mutation accumulation interact to counter TE invasions is key to our understanding of TE expansion dynamics.

In this work, we identify the factors driving the invasion of a highly active transposon in the fungal pathogen *Zymoseptoria tritici*. The pathogen attacks wheat and causes global losses in wheat production (21, 22). The species experienced population-level bursts of TEs linked the creation of genomic diversity and rapid adaptation (23–26). Despite active RIP and RNAi, some *Gypsy* retroelements and class II TIR elements have recently accumulated high copy numbers (24). A substantial number of TEs undergo de-repression when the pathogen colonizes the plant host suggesting that the pathogen lifestyle constitutes risks to effective TE control (12). Recent work has shown that transcription of a retrotransposon named *Styx* negatively impacts asexual reproduction (27). *Styx* was also linked to a large-scale chromosomal rearrangement and copy number variation among field isolates (27). The propensity to create new copies in the recent evolutionary history of the species makes *Styx* transposon an ideal element to investigate how TEs activate and propagate in genomes.

Here, we recapitulate the evolutionary history of the *Styx* element across recent speciation events using reference-quality genomes of the *Zymoseptoria* plant pathogen genus. We focus on *Z. tritici* showing the most recent *Styx* activity. *Styx* copy-number estimates derived from a 900-genome panel reveal two substantial recent expansions in geographically restricted populations. Using four-generation pedigree with completely assembled genomes, we track active transposition of *Styx*. We show that *Styx* triggered several distinct rearrangement types including deletions and duplications of flanking regions, as well as chromosomal fusions. In conjunction, we establish how a recently reactivated TE escapes genomic defenses and triggers genomic rearrangements at observable rates.

## Results

### Evolutionary origins and reactivation across speciation events

The *Styx* transposon is one of the two most active transposons in the fungal wheat pathogen *Z. tritici* shown to recently create new copies and segregate significant copy-number variation within the species (17, 28). *Styx* is restricted to the *Zymoseptoria* genus (27). To recapitulate the evolutionary history of the element, we searched 20 reference-quality genomes defining the global pangenome of *Z. tritici* and four genomes of additional species of the genus (29, 30). *Z. tritici* diverged from its closest sister species *Z. pseudotritici* ∼11,000 years ago (31). *Z. tritici* originated in the Middle East and subsequently colonized North Africa and Europe with the latest spread occurring in the Americas and Oceania following the establishment of wheat cultivation in the past centuries (28, 32). Individual *Zymoseptoria* sp. genomes carry between 0 and 34 *Styx* copies for a total of 200 discovered copies (Figure 1A). The highest number of copies was found in an Argentinian *Z. tritici* isolate (Arg00; 34 copies) followed by the sister species *Z. pseudotritici* and *Z. passerinii* (25 and 23 copies, respectively). Isolates sampled near the center of origin of *Z. tritici* carry no copies of the element (IR01_26b, IR01_48b, ISY92, KE94, TN09, YEQ92). High copy numbers of *Styx* in sister species and the absence of *Styx* in center of origin populations suggests that *Styx* was nearly eliminated from *Z. tritici* early in speciation. *Styx* carries highly conserved TIRs with 97-100% identity overall. Single TIRs (*i.e.* TIRs without a matching partner) are present in all genomes of the genus including in the *Z. tritici* center of origin populations (Figure 1A). Alignment of TIR sequences delineates two groups with one group being exclusive to *Z. passerinii* and one group shared among the other members of the genus. The two TIR groups diverge at a 10-bp motif in the center of the TIR, suggesting that the two TIR groups define independent transposition activity of *Styx* (Figure 1B).

**Figure 1:**
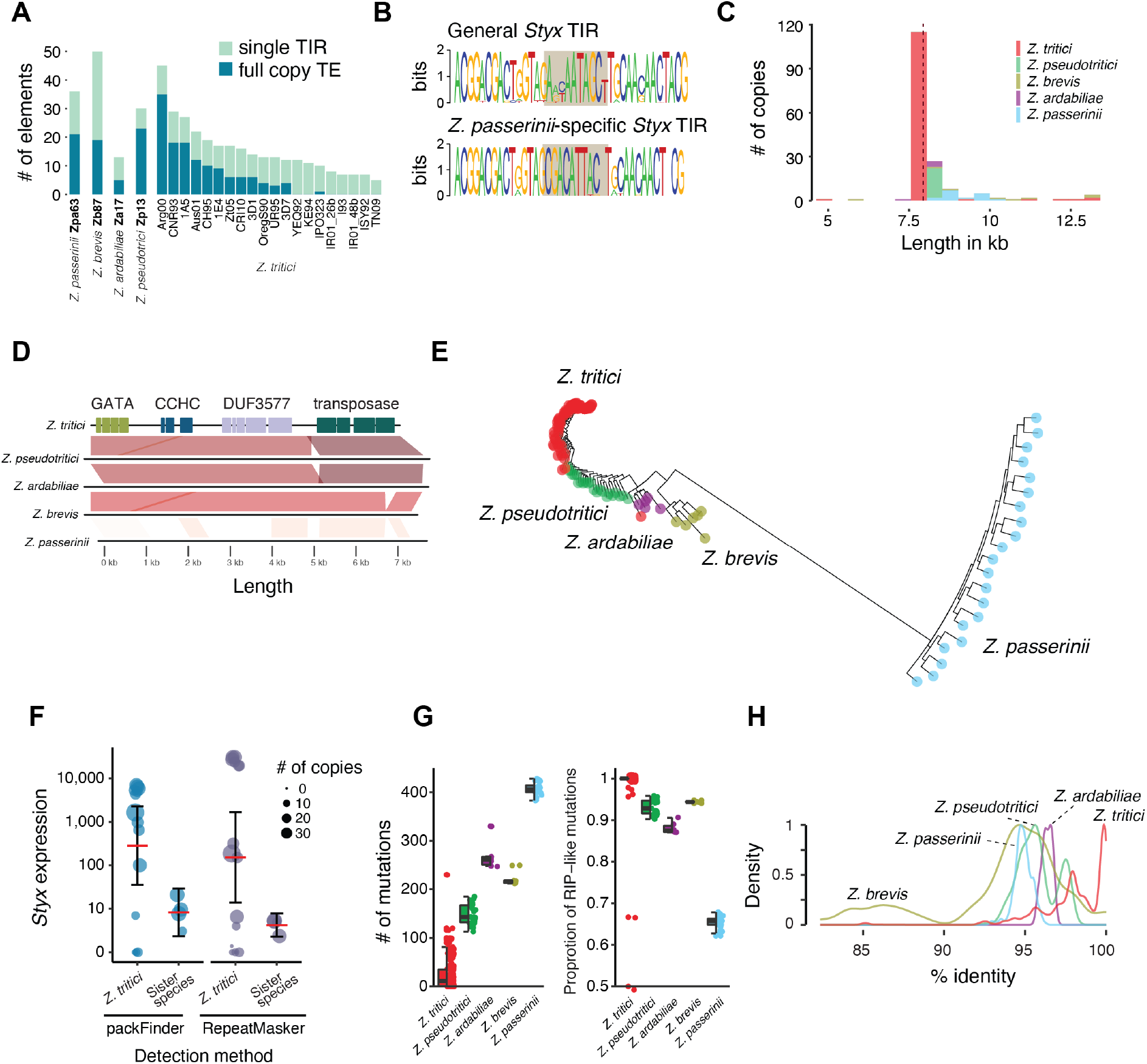
Structure, evolution and expression dynamics of the *Styx* transposable element in the *Zymoseptoria* genus of plant pathogens. **A.** Number of *Styx* copies identified in 24 complete genomes of the *Zymoseptoria* genus. Full elements and single terminal inverted repeats (TIR) are shown separately. **B.** Sequence logo showing the conservation of the TIR sequence identified in *Z. tritici* and *Z. passerinii,* respectively. **C.** Length variation of *Styx* copies across the four *Zymoseptoria* species restricting for copies shorter than 13 kb. **D.** Synteny of the five representative *Styx* sequences from *Z. tritici*, *Z. pseudotritici*, *Z. ardabiliae*, *Z. brevis* and *Z. passerinii* showing syntenic sequences and the insertion present in *Z. pseudotritici*. Darker colors indicate higher sequence identity. **E.** Phylogenetic tree based on the integrase coding sequence of the different *Styx* copies. The tree was rooted using the single degenerate copy present in the *Z. tritici* reference isolate IPO323. **F.** Expression in counts per million of the *Styx* element as annotated by packFinder and RepeatMasker in *Z. tritici* isolates and the four other sister species. Dot size shows the element copy number in the respective isolate. **G.** Number of mutations detected in the integrase coding sequence for all *Styx* copies in the five *Zymoseptoria* species. C->T and G->A transitions are considered RIP-like mutations. **H.** Pairwise identity based on reciprocal blast of the *Styx* copies analyzed separately for each of the five *Zymoseptoria* species.

The majority (∼52%) of *Styx* copies are 7,927 bp in size corresponding to the consensus sequence (Figure 1C). The *Styx* carries four open reading frames, one encoding the putative transposase, two genes with putative DNA binding motifs (GATA and CCHC domains) and a gene with a domain of unknown function (DUF3577) (Figure 1D). Sequences of atypical length are strongly differentiated and likely constitute degenerated copies. The *Z. pseudotritici* genome carries 20 copies ranging from 8,232-8,240 bp in length sharing a unique 285 bp insertion flanked by a tandem repeat of 32 bp suggesting the emergence of a *Styx* sub-family (Figure 1D). Four *Z. ardabiliae* copies carry a similar insertion consistent with a *Styx* sub- family diversification within the genus. The longest *Styx* variant is shared among two species and may constitute a derivative of the ancestral *Styx* present at the origin of the genus. The expansion outside of the *Z. tritici* center of origin was driven by a shorter variant. We performed an ancestral state reconstruction of long versus short *Styx* sequence variants to assess the evolutionary history of *Styx* variants (Figure 1E). The ancestral *Styx* element was most likely shorter lacking the characteristic 285 bp sequence insertion shared by *Z. pseudotritici* and *Z. ardabiliae*. Sequences of the transposase transcript (as defined by the *Z. tritici* consensus) cluster at the species level except for a copy in the genome of the *Z. tritici* strain IPO323, which may represent an ancestral polymorphism shared with *Z. ardabiliae* (Figure 1E). The highest degree of conservation of the transposase locus is found in *Z. tritici* among isolates from Europe, the Americas and Australia with fewer than ten mutations overall. All four open reading frames of the *Styx* are transcribed both on the host and in culture condition in *Z. tritici* (27). Yet, we found the *Styx* element to be largely silenced in culture condition in the sister species (Figure 1E). In contrast with the low expression in these sister species, the high sequence similarity *Styx* copies within *Z. tritici* is coinciding with high levels of transcription (Figure 1E). Consistent with the observed silencing, no sister species *Styx* carries an intact integrase coding sequence.

Copies of the *Z. passerinii Styx* are highly divergent consistent with the accumulation of random mutations (Figure 1G). In contrast, *Z. tritici* copies are highly similar and most divergence was caused by mutations typically generated by RIP genomic defenses (*i.e.* C->T and G->A transitions; Wilcoxon rank sum test *p*-value < 0.05). Around 38% of *Z. tritici Styx* copies retained a pairwise identity ≥99%, consistent with ongoing mobilization of the element and weak effect of RIP (Figure 1H). In *Z. ardabiliae*, *Z. passerinii* and *Z. pseudotritici,* copies share a pairwise identity between 90-99%. In *Z. brevis,* 18% of the *Styx* copies share ≥90% identity and only ∼4% of the copies share ≥99% identity. In conjunction, the degree of sequence conservation, copy number increase and transcriptional activity strongly suggest that *Styx* was recently reactivated in *Z. tritici*.

### Recent *Styx* expansion in South American populations

To recapitulate the temporal and spatial dynamics of the *Styx* expansion in *Z. tritici*, we analyzed a 923-genome panel of isolates collected from wheat fields across the world. To account for limitations of short-read sequenced genomes, we mapped individual sequencing reads to TE consensus sequences to estimate copy numbers (Figure S1). We validated the accuracy of the copy number estimation by analyzing a set of ten isolates with both short read data and completely assembled genomes available (Figure 2A). We find substantial *Styx* copy number variation within the species and across geographic regions (Figure 2B). Genomes sequenced from center of origin populations in the Middle East and North Africa are largely devoid of *Styx* copies. European and North American genomes show an average of 8-9 *Styx* copies per genome, while the Oceanian and South American populations carry on average 17 and 20 copies per genome, respectively (Figure 2B). The highest copy numbers globally were found in genomes from Argentina with a maximum of 56 copies (average of 31 copies). In contrast to strong geographic effects worldwide on *Styx* copy numbers, copy numbers were largely stable across the sampling period 1999-2016 in Europe with overall moderate levels of *Styx* (Figure 2C).

**Figure 2:**
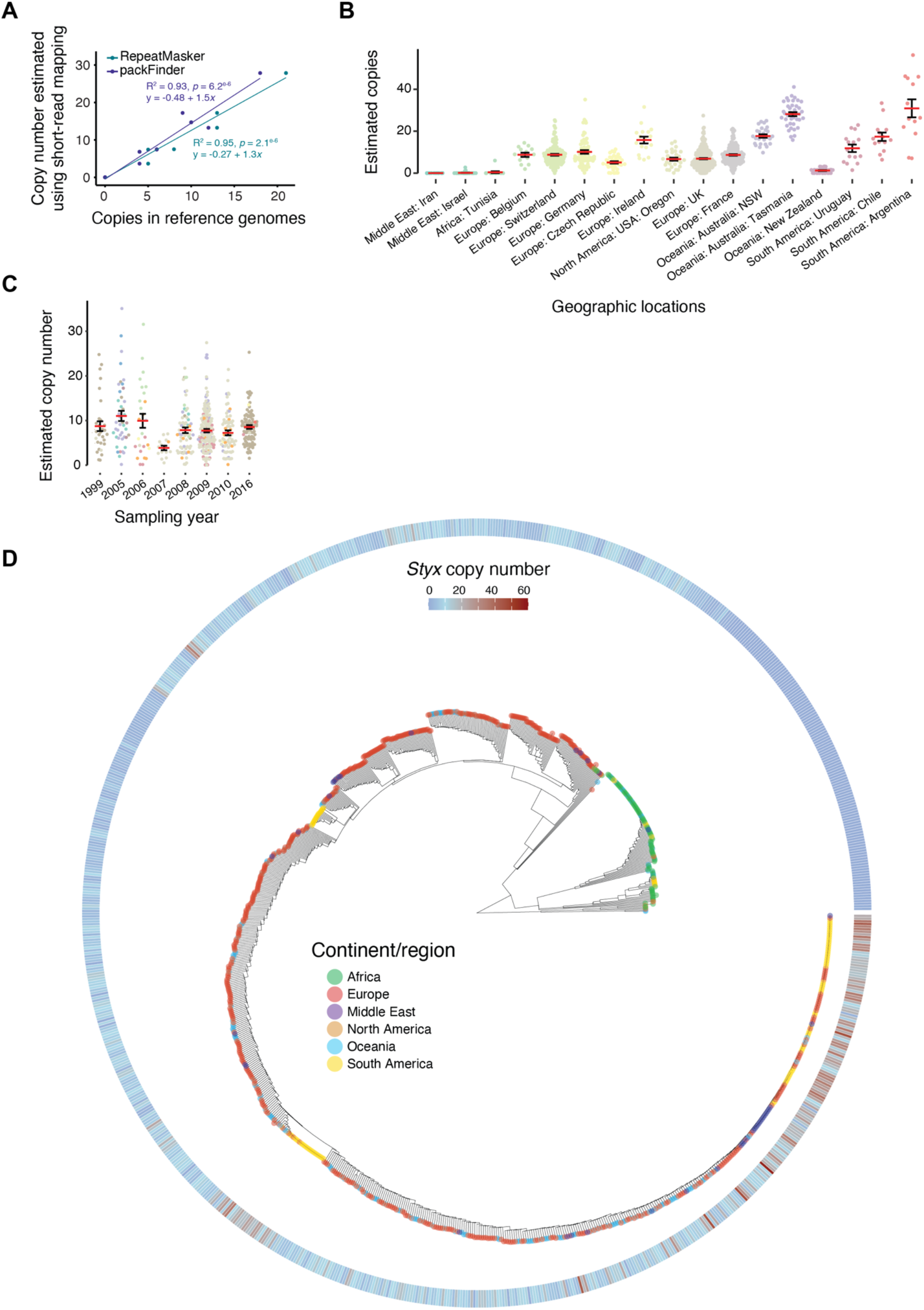
Reactivation of the *Styx* element in the wheat pathogen *Z. tritici* tracked by a 900-genome panel. **A.** Correlation between the number of annotated *Styx* copies from RepeatMasker or packFinder based annotation of chromosome-level assemblies and the estimated copy number from short-read mapping on the consensus sequence of the *Styx* element. **B.** Estimated *Styx* copy number in *Z. tritici* isolates sampled across the species geographic range (17 geographic locations with more than 10 isolates for a total of 856 isolates). **C.** Estimated *Styx* copy number in isolates sampled in Europe between the years 1999 and 2016. **D.** Phylogenetic tree based on SNPs identified against the *Styx* consensus sequence in 923 worldwide *Z. tritici* isolates. The heatmap on the outside of the tree shows the estimated *Styx* copy number per sequenced genome.

Newly created copies during the *Styx* expansion may have started to accumulate mutations. Based on the 923-genome panel, we identified a total of 1,707 single nucleotide variants affecting 21% of the 7,927 bp *Styx* consensus sequence. Consistent with the high copy-numbers and recent expansion, *Styx* copies in Oceanian and South American populations were highly similar (Figure 2D). Center of origin populations tend to carry more sequence variants consistent with the presence of older *Styx* copies. Isolates from Africa carry an average of 447 variants per copy, which is consistently higher than in other regions (Tukey’s HSD, *p*-value < 0.01; Figure 3A). Mutations in *Styx* affected primarily non-coding regions (∼43%; Figure 3B). An additional 30.5% and 4.5% of the mutations were missense and stop gain variants, respectively. C->T and G->A transitions dominate the mutation spectrum consistent with RIP genomic defenses driving the divergence of *Styx* copies (Figure 3C). *Styx* copies in Oceanian genomes have a higher proportion of frameshift and missense variants despite overall lower differentiation among copies (Tukey’s HSD; *p*-value < 0.001; Figure 3D). The broad genomic survey of the species reveals a strong association of rapid *Styx* copy number expansion following founder events at the origin of Oceania and South America populations, as well as a concurrent degradation of genomic defenses against *Styx*.

**Figure 3:**
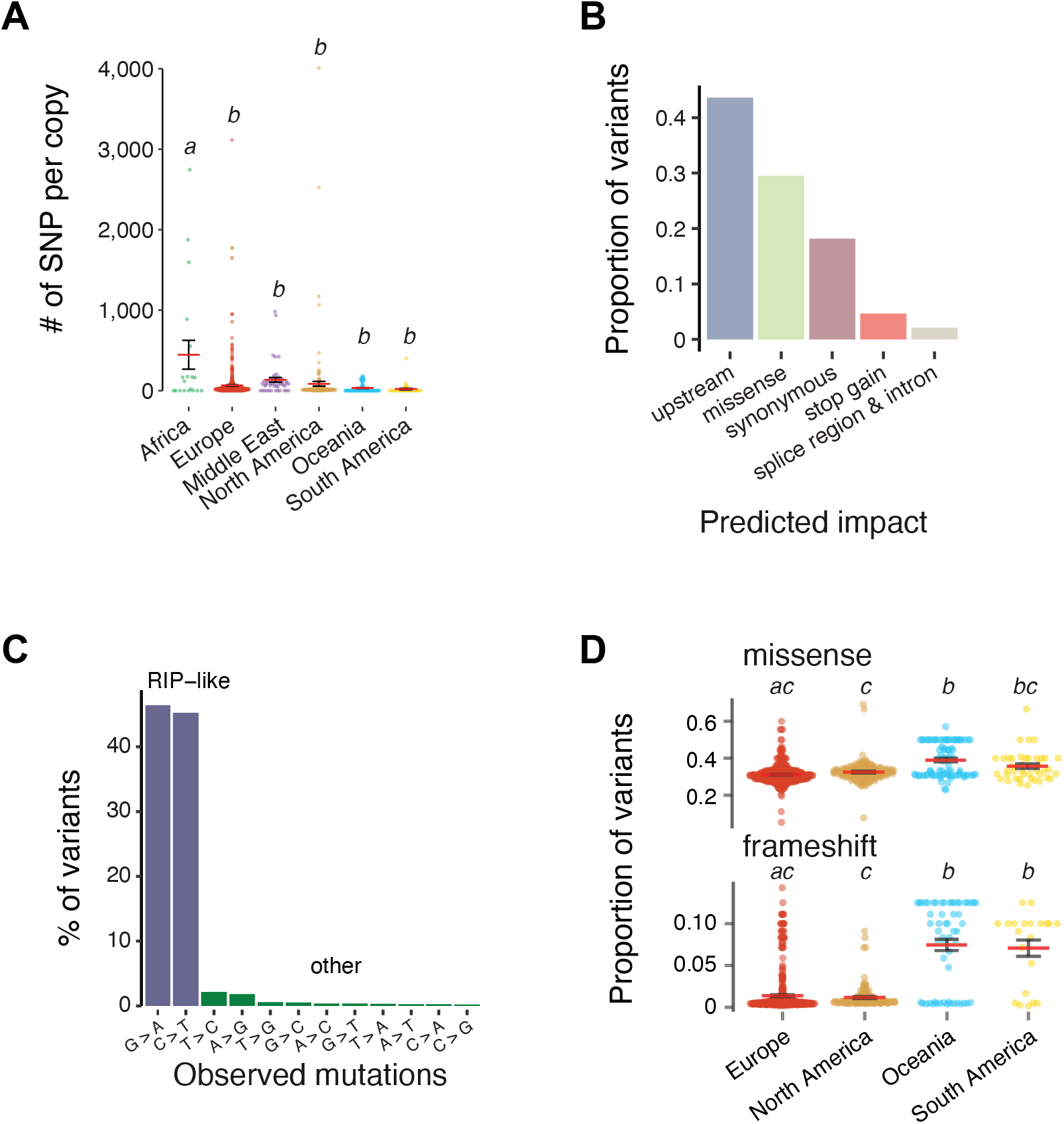
Mutation accumulation of the *Styx* element in *Z. tritici* driven by genomic defenses. **A.** Average number of variants detected in reads from each of the 923 worldwide *Z. tritici* isolates mapped against the *Styx* consensus sequence and normalized by the estimated *Styx* copy number. **B.** Predicted effect of variants detected in the read alignment among all isolates. **C.** Proportion of biallelic variants classified according to the type of transition or transversion. C->T and G->A transitions are considered RIP-like mutations. **D.** Proportion of variants annotated as missense and frameshift. Letters indicate significant differences (Tukey HSD, *p* < 0.05).

### Styx mobilization and chromosome-scale rearrangements upon meiosis

Highly active TEs such as the *Styx* element may show observable rates of transposition even over single generations. Investigating the youngest insertions of *Styx* provides a more complete spectrum of insertion sites as selection may not have eliminated yet all deleterious new insertions. We analyzed evidence for newly inserted copies of *Styx* in a four-generation pedigree initiated by two isolates collected from a European wheat field (1A5 and 1E4, (33). The pedigree comprised a total of 10 progeny and revealed a large rearrangement of chromosome 17 most likely triggered by copies of *Styx* (Figure 4A, (34). The parents 1A5 and 1E4 carry 18 and 9 *Styx* copies, respectively, and progeny carry 9-16 copies (Figure 4B). We analyzed recombination tracks using whole-genome alignments of each progeny genome against the two parental genomes to differentiate transposition creating a new *Styx* copy from recombination introducing existing *Styx* copies into a new background. Using this recombination map established for each progeny, we tracked vertical inheritance of each *Styx* copy in the pedigree (Figure 4C). We found that both parental genomes contributed equally to copy numbers in the progeny with an average of 3.8 and 4 copies originating from the 1A5 and 1E4 parent, respectively. A further 44 *Styx* copies (∼36 %) from all progenies were found in chromosomal locations incompatible with vertical inheritance from either of the two parental backgrounds and, hence, constitute new insertions (Figure 4D).

**Figure 4:**
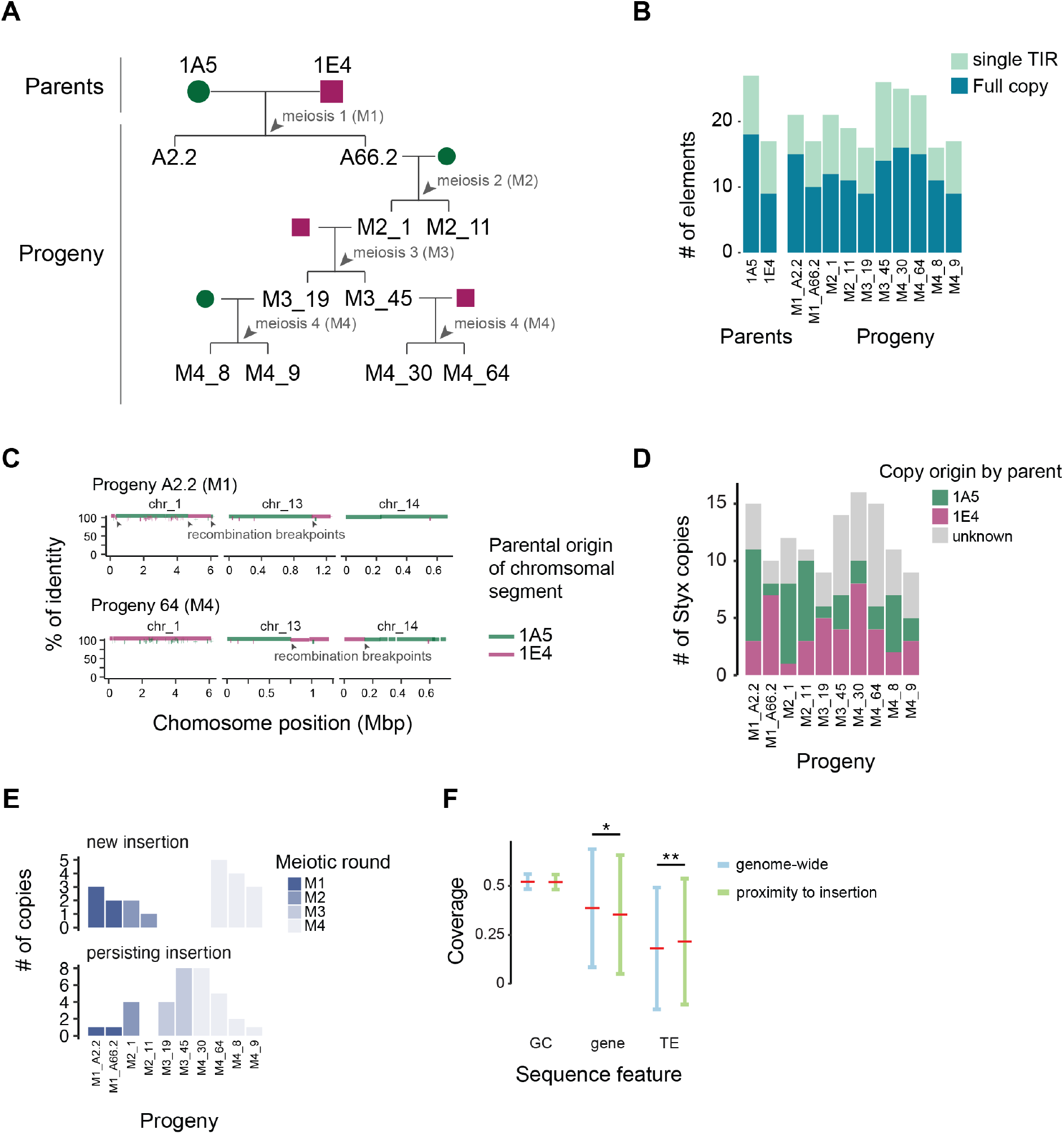
Four-generation pedigree analyses reveals active transposition of the *Styx* element. **A.** Schematic of the analyzed *Z. tritici* pedigree with parent-offspring relations. **B.** Number of *Styx* copies identified in the pedigree. Full elements and single terminal inverted repeats (TIR) are shown separately. **C.** Illustration of the chromosomal block synteny analyses of the progeny isolates M1_A2.2 and M4_64 showing evidence for recombination events on core chromosomes 1 and 13 and accessory chromosome 14. **D.** Parental origin of the *Styx* copies in different progeny isolates. **E.** Status of the *Styx* copies identified in each meiotic round along the progeny (M1 to M4). **F.** Isochore, gene and transposon coverage in proximity to *Styx* insertions (windows of 50 kb centered on *Styx* copies) compared to a genome-wide set of 50 kb windows. Asterisks indicate the result of a pairwise Wilcoxon rank sum test (* < 0.05, ** < 0.01 after Holm correction).

We searched for ancestral sequences at the origin of each new *Styx* insertion including the 6- bp target site duplication sites. We discarded loci where parental genomes segregated presence-absence of *Styx* copies. Each progeny genome carries between 1-10 new *Styx* insertions for a total of 32 distinct insertion events (Figure 4E). New insertions cover 7,927+/-1 bp of sequence matching the full-length *Styx* element and locate to 8 out of 13 core chromosomes, as well as 5 out of 8 accessory chromosomes. New insertions occurred in regions with higher TE density compared to the genome-wide average (Figure 4F; Wilcoxon rank sum test *p*-value < 1e-3). New insertions tend to persist in the genomes as the number of inserted copies increases but are rapidly lost further down the pedigree (*i.e.*, meiotic round 3 to 4) consistent with recombination reducing *Styx* copies in progeny genomes (Figure 4E).

We more closely analyzed the association between recombination breakpoints and *Styx* copies. We identified a total of 138 recombination breakpoints in the 10 progeny genomes with an average of ∼14 breakpoints per progeny (Figure 5A). We assessed co-occurrence in 50-kb windows and found that *Styx* copies were over two-fold enriched near recombination breakpoints (Fisher’s exact test odds ratio = 2.4, *p*-value = 0.03). Genomic features such as isochores can co-vary with recombination rates (35, 36). In the pedigree, recombination breakpoints occurred predominantly in gene-rich and TE-poor regions sharing similarities with the main genomic niche of *Styx* (Tukey’s HSD; *p*-value < 1e-16; Figure 5B). The most striking association of *Styx* copies with recombination breakpoints is a large chromosomal translocation involving core chromosomes 6 and 12 (Figure 5C). The two *Styx* copies inherited from the 1A5 parental genome were near the synteny breakpoints at the origin of the fused core chromosomes in progeny M4_8 (Figure 5C).

**Figure 5:**
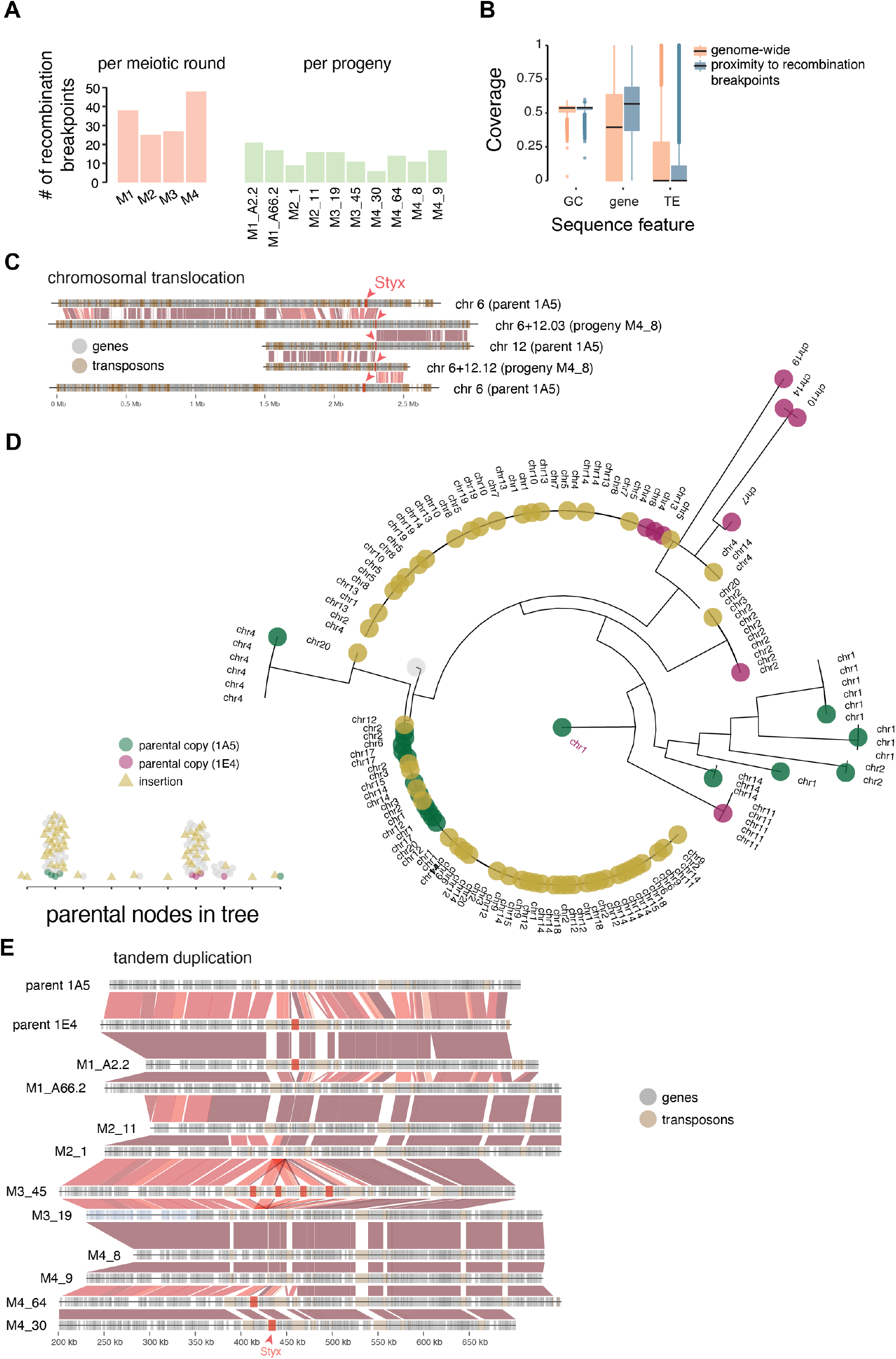
Chromosomal translocations triggered by new *Styx* insertions and insertion preference near recombination breakpoints. **A.** Number of recombination breakpoints identified in the progeny genomes. Numbers are shown per meiotic round (M1 to M4) and separately for each progeny isolate. **B.** Isochore, gene and TE coverage at recombination breakpoints compared to genome-wide 50 kb windows. **C.** Synteny plot illustrating the chromosomal translocation that occurred between chromosomes 6 and 12 in the progeny isolate M4_8. *Styx* copies are highlighted with red arrows. **D.** Phylogenetic tree based on full *Styx* sequences detected in parental and progeny genomes. Vertically inherited *Styx* copies are differentiated by colors from new insertions. **E.** Synteny plot illustrating the *Styx* tandem duplication that occurred on chromosome 8 in the progeny isolate M3_45. *Styx* copies are highlighted in red, other TEs in brown and genes in blue.

The recency of the new *Styx* insertions enables to track the genealogical history of copies across pedigree generations. For this, we performed full-length multiple sequence alignments of all *Styx* copies detected among parental isolates and progeny (Figure 5D). The 32 new *Styx* insertions separate into two distinct clades with one composed of 21 sequences being nearly identical to 11 copies of *Styx* identified in the parental genome 1A5. A second clade is composed of eleven insertions nearly identical to three *Styx* found in the parental genome 1E4 (Figure 5D). We retraced the origin of a new *Styx* insertion on chromosome 1 found in progeny M2_1 (position 3,640,021 bp and target site duplication ATCGAG) to a 1A5 parental copy on chromosome 2 (position 52,220 bp and with target site duplication CGTGAT). We also retraced a second insertion on chromosome 3 found in the progeny M4_64 (position 1,432,987 bp with target site duplication TGTGAT) to a 1E4 parental copy on chromosome 2 (position 3,217,078 bp with target site duplication CATAAG; Figure 5D). Interestingly, the 1E4 parental copy encodes a non-functional integrase coding sequence indicating that even degenerate copies of the *Styx* can be mobilized by an integrase protein encoded by intact copies. Two *Styx* insertions each on chromosomes 14 and 15 were recurrent insertion events within 100 kb distance from each other (Figure S2). Overall, 44% (8 out of 18) of the new insertions on the same chromosome occurred within 150 kb from the parental copy. We observed a similar transposition pattern of *Styx* copies across *Zymoseptoria* species. Overall ∼51% of *Styx* copies are clustered on the same chromosome and separated by less than 150kb from each other. Physical proximity of new *Styx* insertions suggests that chromosomal conformation (*i.e.* contact) may be relevant to the transposition of the element.

We identified a total of 25 solo-TIRs among progeny, which is comparable to the abundance of solo-TIRs across *Zymoseptoria* species (Figure 4B). However, solo-TIRs are readily created *de novo* as identified in the progeny isolate M3_45 with three new solo-TIRs compared to parental genomes. Analyzing synteny at the corresponding locus on chromosome 8 showed that the three solo-TIRs originated from a triplication of a *Styx* element in the parent 1E4 (Figure 5E). The parental *Styx* duplicated and inserted three times near the original location creating 2-3 additional copies of six genes near the original *Styx* element. The six genes include a duplicated secreted endonuclease gene, a triplicated gene encoding a secreted protein of unknown function, a twice duplicated gene encoding a membrane-bound protein, a duplicated gene encoding a predicted protein kinase and two genes encoding for proteins of unknown function (duplicated and triplicated, respectively). Across *Zymoseptoria* species, we find ∼19% of all *Styx* copies within 50 kb of two or more gene duplicates (23 of 122 total copies). Similarly, we found tandem-inverted *Styx* copies co-localized on the same chromosome arm. Hence, the *Styx* transposition may be driven by proximal insertions and topological preferences (Figure S3).

Given the ability of the *Styx* element to mobilize over single rounds of meiosis and trigger sequence rearrangements, we investigated whether genomic defenses counteracted *Styx* activity. The average number of *Styx* copies remained largely stable in the pedigree despite evidence for new insertions (Figure 4B-E). Hence, either strong purifying selection eliminated progeny with higher *Styx* copy numbers or *Styx* copies were readily excised. We find evidence for excision at a *Styx* locus of the 1A5 parental chromosome 10, which is missing *Styx* in two progeny isolates. The excision event is further supported by the retention of parental sequences surrounding the element (Figure S4). Synteny analyses of the excision locus revealed three paralogous genes on either side of the parental *Styx*. The duplicated genes in the flanking region likely served as homology anchors for the excision of the *Styx* copy (Figure S4). Taken together, *Styx* mobilization is likely an agent triggering chromosomal rearrangements over single generations and impacts adjacent chromosomal regions by possibly favoring gene duplications.

### *Styx* activity escapes genomic defenses by the RIP machinery

Repeat-induced point (RIP) mutations is a mechanism that inactivates TEs in fungal genomes. Mutational signatures of active RIP were found across the *Zymoseptoria* genus (37). Here, we analyzed *Styx* copies in the progeny to quantify the occurrence of *de novo* RIP-like mutations. We focused on the 31 *Styx* copies that persisted for one to four rounds of meiosis and identified mutations in new copies using the parental copy of the element as a reference. Over 93% of the *Styx* copies remained identical throughout the pedigree (29/31). The two *Styx* copies with point mutations have 5+12 and 9+10 RIP-like mutations respectively (*i.e.*, C->T and G->A transitions). None of the new *Styx* copies in the pedigree showed signatures of RIP. To address whether RIP is broadly ineffective against TEs, we investigated the accumulation of *de novo* mutations in four additional TEs with varying copy numbers and expression profiles (RLC_Deimos, DXX_Birute, DTA_Vera and DTT_Tapputi). Identical to the procedure used to recover *Styx* copies, we analyzed target site duplication sequences to recover orthologous copies among pedigree genomes. The DTA_Vera and DTT_Tapputi elements have 12 and 14 copies in the pedigree, respectively, although copies share only low sequence identity and were likely not targeted by RIP (0.55 and 0.43 maximal pairwise sequence identity, respectively). For the elements DXX_Birute and RLC_Deimos, *de novo* mutation accumulation was only detected in the high-copy RLC_*Deimos* (74 copies in the 1A5 parent) despite copies of the element sharing between 0.4 and 0.9 sequence identity. The 21 detected mutations were restricted to 8 out of 142 copies and were only found in progeny genomes of the first round of meiosis (*i.e.*, M1_A66.2 and M1.A2.2). Given the older sequencing technology used to generate PacBio long-reads (RSII vs. Sequel), we cannot rule out that non-polished sequencing errors persisted in the assembly as artefacts. Altogether, we conclude that genomic defenses in *Z. tritici*, including RIP, are unable to efficiently target *Styx* or other TE copies. The rapid expansion of the *Styx* transposon in the pedigree is consistent with the expansion dynamics observed at the species level with major consequences for chromosomal integrity and the faithful transmission of genetic information.

## Discussion

### A recent origin of Styx within the Zymoseptoria genus

TEs are major drivers of genome evolution over deep time scales. Here, we unravel the recent evolutionary history of the *Styx* transposon and how this recapitulates major phases of TE family dynamics. The *Styx* recently differentiated into sub-elements with distinct TIRs and sequence length concomitant with speciation of five closely related *Zymoseptoria* species. Consistent with a recent origin, two of the four *Styx* coding sequences are exclusively found in *Zymoseptoria* species (27). The encoded DDD/E transposase suggests a common origin with members of the IS3EU DNA transposon superfamily (38). TE families typically differentiate rapidly with few documented examples of ongoing activity such as the high-copy miniature non-autonomous *mPing* and the two autonomous *Ping* and *Pong* elements in rice retaining signatures of a common ancestor (19). Given the shared coding sequences with IS3EU family members, *Styx* likely evolved through gene co-option in the ancestor of the five *Zymoseptoria* species. The IS3EU transposase family shares similarities with bacterial insertion sequences from the IS3 family (38, 39). Furthermore, horizontal transfer of DDD/E TE families is widespread and found in multiple kingdoms (40, 41). However, no *Styx*-like elements were found in species outside of the *Zymoseptoria* genus.

### Independent waves of *Styx* reactivation

Sequence identity patterns of *Styx* copies in the *Zymoseptoria* genus are consistent with at least two recent independent reactivation events in the ancestors of *Z. pseudotritici* and *Z. tritici* (Figure 6). In *Z. tritici*, high-quality genomes analyzed across the global distribution range carry highly heterogeneous copy-numbers suggesting very recent transposition activity (17). In concordance, we find that the element is highly expressed in many *Z. tritici* isolates and silenced in other species of the genus indicating that copies were repressed following the transposition bursts in these species (Figure 6). Host control over TEs can be mediated by mutations affecting transposon mobility (18, 19, 42). Here, we investigated proximal mechanisms associated with *Styx* mobilization and repression. The TIRs of the *Z. passerinii Styx* sub-element are mutated compared to the dominant TIR sequence found in the genus likely affecting *Styx* mobility. Similarly, the recent *mPing* burst in rice was likely facilitated by a point mutation adjacent to the *mPing* TIRs (J. Chen et al., 2019). How transposons reactivate over evolutionary time scales remains largely unknown. However, external stressors such as temperature can promote transposon activity (11, 16). In *Z. tritici,* nutrient limitation and stress imposed by plant immune defense favor de-repression in a TE-specific manner (12). It remains unknown whether resistant wheat cultivars and increased fungicide applications are constituting sufficiently strong factors to cause lasting TE de-repression in *Z. tritici*. Fungicide exposure under laboratory conditions can lead to transposon activity in other pathogens (43).

**Figure 6:**
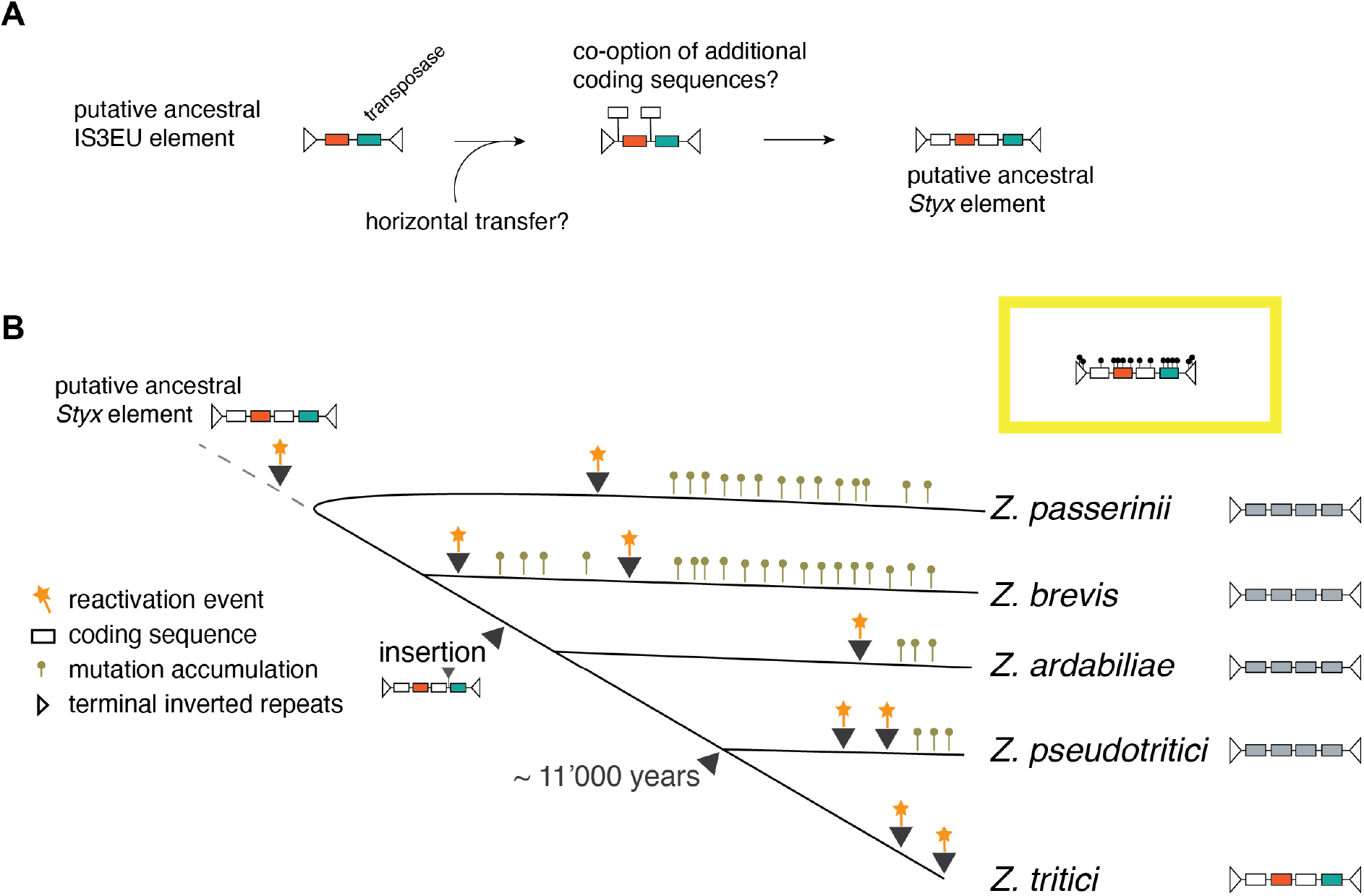
Proposed model illustrating the sequence of evolutionary events at the origin of the multiple *Styx* expansions in the *Zymoseptoria* genus. A. *Styx* likely originated from an ancestral class II IS3EU element encoding two proteins including a DDD/E transposase required for mobilization. After a putative horizontal transfer in an ancestor to the *Zymoseptoria* genus, the element co-opted two additional coding sequences to form the *Styx* element. B. The newly formed *Styx* element possible activated first prior to speciation in the genus. Following speciation, *Styx* experienced substantial sequence divergence. Based on species-level copy identity, *Styx* likely reactivated independently two times in *Z. brevis*, *Z. pseudotritici* and *Z. tritici.* In *Z. tritici*, the mobilization likely occurred following the spread of the pathogen outside of the center of origin in the Middle East.

The genome-wide transposon content in *Z. brevis and Z. passerinii* is higher than in *Z. tritici* consistent with more young elements and recent activity (37). However, the same two species have stronger signatures of RIP-like mutations across their TE repertoire suggesting that higher TE activity is counterbalanced by TE defenses. In agreement with RIP being active in the genus, a large fraction of the mutations identified across *Styx* copies were most likely generated by TE defenses. Low activity of RIP in *Z. tritici* is most likely explained by the near complete loss of a DNA methyltransferase known to promote cytosine methylation and RIP in *Neurospora crassa* (44–46). We show therefore a parallel between *Styx* copy number and the presence of *DIM2* in the genus that suggests that RIP might contribute to *Styx* control.

### A population-level perspective on TE reactivation

*Styx* copy number estimates across >900 *Z. tritici* genomes vary by an order of magnitude across the global distribution range. The element is nearly absent from the center of origin populations in the Middle East and North Africa. *Styx* copies culminate in populations from Oceania and South America suggesting a recent reactivation in these regions. In agreement, we find that *Styx* copies from Oceania and South America accumulated very few mutations and copies retained high sequence identity. The transposon content of genomes is expected to result from the equilibrium established between transposon activity, host repressive mechanisms and purifying selection (1). However, bursts in transposon activity and population bottlenecks following founder events are impacting transposon dynamics as well (47). The evolutionary history of *Z. tritici* is tightly linked to wheat domestication in the Fertile Crescent followed by stepwise introduction events to new continents (28). Populations outside of the Middle East and Europe have both gained in transposon content and overall reduced RIP-like mutations (28). Colonization events over the past centuries in Australia induced population bottlenecks leading to reduced genetic diversity and likely less efficient selection. Such relaxation in selection may have weakened genomic defenses against transposons. Relaxation of genomic defenses likely underpins also incipient genome size expansion within the species (24). *Styx* follows the same trend of higher copies numbers in regions with weakened genomic defenses (*i.e.* RIP-like mutations). The proximal trigger of *Styx* reactivation outside of the *Z. tritici* center of origin may well have been the loss of *DIM2* underpinning RIP activity.

### *Styx* escapes genomic defenses and destabilizes chromosomal sequences

How transposons activate and reintegrate into chromosomal sequences remains poorly understood. A particular challenge arises from the action of purifying selection removing an unknown subset of all insertions. In addition, population bottlenecks and founder events can arbitrarily eliminate insertions from populations. Here, we used a pedigree to track the *Styx* element over generations using complete genome sequences. *Styx* is indeed mobile creating dozens of new copies through a copy-and-paste replication mechanism. Retrotransposon-like copy-and-paste transposition is consistent with the substantial gains in copy numbers observed across the species range. By tracking new copies to their progeniture sequence in parental genomes, we found that both copies with functional and degenerate transposase coding sequences are mobile. Degenerate copies likely benefited from transposase expression of intact copies. Mobility retention of degenerate *Styx* may favor even further sequence reduction to reduce into miniature inverted-repeat TEs as shown for the *mPing* elements (7, 19).

The new *Styx* insertions in the *Z. tritici* progeny are found preferentially closer to genes. However, *Styx* shows no detectable insertion site sequence preference suggesting that the heterogeneous distribution in the genome across the species most likely arises from preferences for open chromatin but may also be influenced by random factors such as genetic drift. In the pedigree, *Styx* loci display precise excisions mediated by recombination of flanking sets of paralogous genes. Furthermore, recombination breakpoints are preferred *Styx* insertion sites in the pedigree. Associations of TEs and recombination breakpoints were previously observed for *Drosophila P*-elements inducing recombination in males and promoting large deletions and duplications of flanking regions (48). TE-mediated recombination such as driven by the RAG recombinase system evolved from transposon domestication (49). Reshuffling of vertebrate immune loci is mediated by the RAG recombinase system with suppressed transposition activity.

Some of the most consequential effects of unchecked TE activity in genomes are large sequence rearrangements. CACTA-elements for instance are associated with chromosomal rearrangements that led to the formation of the R-r complex that controls the production of anthocyanin in maize (50). LINE-1 insertions have been associated with large deletions and the amplification of oncogenes in cancer cells (51). The *Styx* likely contributed to multiple large chromosomal rearrangements over only four generations and a few dozen new insertions overall. The chromosomal fusions associated with *Styx* activity have the potential to initiate chromosomal sequence meltdowns. Deleterious effects of *Styx* are likely compounded by the tendency to generate paralogous gene copies. This is consistent with double-strand breaks being involved in the *Styx* replicative-transposition. Double-strand breaks would have favored the observed tandem duplications of the element and concurrent gene duplications in flanking regions. Paralogues in proximity to *Styx* insertion sites are particularly noteworthy given the very low rate of gene duplicate retention compared to other fungi likely as a consequence of active RIP (29, 52). Taken together, *Styx* plays complex roles in the *Zymoseptoria* genus from influencing asexual reproduction, contributing to gene duplications, chromosomal rearrangements and fusions. The loss of control over *Styx* transposition in recently established populations highlights the tenuous and transitory control host genomes exert over selfish elements.

## Methods

### Transposable element annotation and insertion genotyping

We accessed previously published complete genome assemblies of 20 reference-quality genomes of *Z. tritici* spanning the global distribution range of the pathogen (29, 30) (Supplementary Table S1). In addition, we analyzed four complete genomes of sister species *Z. passerinii, Z. ardabiliae, Z. brevis and Z. pseudotritici* (30) (Supplementary Table S1). We also analyzed ten complete genomes assembled from a four-generation pedigree started by the isolates 1A5 and 1E4 (34) (Supplementary Table S1). We annotated all genomes for TEs using the method and consensus sequences described in Badet et al (2020). In addition, for *Styx,* we implemented an additional annotation based on the 36 bp TIR as described in Wang et al (2021) to identify conserved copies across the 34 genome assemblies using the R package *packFinder* version 1.6.0 (53). We performed the *packSearch* in R version 4.1.3 using the following parameters: *tsdMismatch* = 6, *mismatch* = 8, *elementLength* = c(100, 30000), *tsdLength* = 6. TIR positions were deduced using the *identifyTirMatches* function of *packFinder*. Single TIR with no co-occurring pair within 30 kb distance were considered solo-TIRs.

For the TEs DTA_Vera, DTT_Tapputi and DXX_Birute, we first performed self-alignments of their consensus sequences to identify TIRs using blastn with *-word_size 4 -perc_identity 85 -dust no* as parameters (fasta sequences are provided in Supplementary Data S1). The TIR was then used to retrieve copies of each element in the parent and progeny genomes of the pedigree using packFinder with parameters adjusted to the element length (*mismatch* = round(*TIR_length* * 0.15), *tsdMismatch* = round(*TSD_length* * 0.4), *elementLength* = c(*TE_length* - 5, *TE_length* + 5) and *tsdLength* = c(2:6)). Identified *Styx* loci and target site duplications were then used to identify homologous positions among *Styx* copies in genomes of the pedigree. For the high-copy TE RLC_Deimos we used RepeatMasker to annotate progeny genomes retaining only annotations covering ≥95% of the element consensus sequence tolerating elements exceeding length by ≤5% (29). To identify copies at homologous positions, we considered five additional base pairs on each side of the Deimos sequences as putative TSD.

To investigate genomic characteristics near TE loci, we binned each genome of the pedigree into 10,000 bp windows to compute gene, TE and GC coverage. We used gene and TE annotations from the Badet et al. (2020) study together with the bedtools *coverage* function.

Similarly, we used the bedtools *nuc* function to compute GC content across all genomic 10 kb windows.

### Sequence alignments and phylogenetic tree inference

All sequence alignments were performed with the software MAFFT v 7.475 using the -- maxiterate 1000 and –leavegappyregion parameters (54). The TIR sequences were recovered based on the positions provided by the *identifyTirMatches* function of the *packFinder* R package and aligned using MAFFT. Sequence logos were calculated using the Berkeley WebLogo server for the aligned TIR sequences from *Z. passerinii* and *Z. tritici* separately (55). For phylogenetic reconstruction, all detected copies of *Styx* across the five *Zymoseptoria* species were aligned jointly with either the *Styx* consensus sequence or the coding sequence of the integrase locus. The resulting alignments were trimmed using the *remove_reference_gaps_in_alignment.pl* script (https://github.com/lakras/bio-helper-scripts/blob/main/aligned-fasta) to the consensus sequence or the integrase gene. Phylogenetic trees were then inferred with FastTree v2.1.11_1 software using 1,000 bootstraps and the generalized time-reversible model (-nt -boot 1000 -seed 1253 -gtr options) (56). Variable positions in the sequence alignments were identified in the trimmed alignments using the *snp- sites* v2.5.1 software (57). Resulting vcf files were simplified to table formats using the *VcfSimplify.py* script (https://github.com/everestial/VCF-Simplify). Bi-allelic SNPs with a C or G as reference allele and T or A as alternative allele, respectively, were considered RIP-like mutations. Tabular data were analyzed in R version 4.1.2 and visualized using the *ggplot2* v3.3.5 package (58).

### Identification of recombination blocks in the pedigree

To recapitulate recombination events across the pedigree, we assigned chromosomal blocks in each of the ten progeny genome to the parent of origin (1A5 or 1E4) using pairwise whole-genome alignments with *nucmer* (MUMmer version 4.0.0rc1) (59). For the alignment step, we used a minimum length of a single exact match of 100 bp, a maximum gap between two matches in a cluster of 10 bp, a minimum cluster of matches of 1000 and an alignment extension in poorly scoring regions of 200 bp (options -l 100 -g 10 -c 1000 -b 200). We filtered the resulting alignments for a minimum identity of 99% and a minimum length of 50,000 bp using *delta-filter* (options -i 99 -l 50000). The filtered alignment coordinates were then converted into a tab-delimited format with the *show-coords* tool (-THrd options). To assign parental origins of progeny chromosomal blocks, we concatenated filtered match coordinates for both parental genomes. We assigned the parent of origin for a chromosomal block based the presence of matches satisfying the >50,000 bp alignment length and ≥99% identity criteria. To assign the parental origin of individual *Styx* copies in progeny genomes, we matched the chromosomal coordinates of *Styx* copies with the chromosomal block information on the parent of origin using *bedtools intersect* version v2.30.0 (60). Shifts in homology from one parent to the other along progeny chromosomes were considered as the most likely recombination breakpoints. To reduce false positives, alignment blocks where both parents showed >99.8 percentage identity and one of the two parental alignment overlapped by more than 50% of their length were left out. Aligned blocks were only retained if the length exceeded 100 kb. Finally, predicted recombination breakpoints separated by less than 20 kb were merged into a single recombination event given that closely spaced recombination breakpoints are unlikely due to crossover interference (61).

### *Styx* expression analysis

We investigated *Styx* expression in the *Zymoseptoria* genus using RNA-seq data generated in axenic growth conditions. Datasets from 17 *Z. tritici* isolates were downloaded from the NCBI BioProject PRJNA559981. In addition, we used Sequence Read Archive (SRA) RNA-seq experiment accessions SRX4341756, SRX4341752 and SRX4341751 for isolate 1E4 and SRX4341748, SRX4341749 and SRX4341750 for isolate 1A5. For *Z. pseudotritici*, *Z. ardabiliae*, *Z. brevis* and *Z. passerinii,* RNA-seq datasets were retrieved from the NCBI BioProjects PRJNA277173, PRJNA277174, PRJNA277175 and PRJNA639021 respectively. The raw reads were trimmed using Trimmomatic v0.39 and mapped to the genome matching the source of the data using STAR v2.7.10a/2.7.9a while allowing for multiple mapped reads (-*outFilterMultiNmax* 100 -*winAnchorMultimapNmax* 200 parameters) (62, 63). Gene and transposon family expression were assessed using TEtranscripts v2.2.1 (-*mode* multi) (64). For downstream analyses, read counts were normalized to counts per million of reads (cpm) in R using the *calcNormFactors* function from the *edgeR* package v3.34.1 (method = “TMM”) (65).

### Estimation of *Styx* copy numbers in genome resequencing datasets

To assess the geographic and temporal variation in *Styx* copy numbers in *Z. tritici*, we used a 923-genome panel of Illumina sequenced genomes (28). Accession numbers for the NCBI SRA are available from Feurtey et al, 2023. The collection covers 42 countries and all continents where the pathogen has been recorded. We trimmed raw Illumina reads to remove adapters and retain only high quality bases (LEADING:15 TRAILING:15 SLIDINGWINDOW:5:15 MINLEN:50) with Trimmomatic v.0.39 (62). We used the *Styx* consensus sequence (29) as a reference sequence to map reads (as single reads ignoring read pair information) with the option --very-sensitive-local using bowtie2 v.2.4.1 (66). Read alignments per analyzed genome were used to derive the number of reads aligning to the *Styx* consensus with the *idxstats* option of *samtools* v.1.10 (67).

