## Supplementary Data for "Recent reactivation of a pathogenicity-associated transposable element triggers major chromosomal rearrangements in a fungal wheat pathogen"

**Data S1**: Sequences of the Terminal Inverted Repeats used for transposon re-annotation using *packfinder.*

>DXX_Styx

ACGGACGACTGGTAGAACAATAGCTGCAAGAACTCG

>DTA_Vera

CAGTCTGCTAAACAATCAAT

>DTT_Tapputi

AGAAGTCACTTGCAAGGTATGCACGGTGCACGCTGTGCACGGTCGCA

>DXX_Birute

ACTCGTGTTGCAAGCTTCTCACCGCCAAAGCGGTGTTGGCCGGCTCAATGAGAATCACCAGCTCAACACGACTGTATC
