## Supplementary Figures for "Recent reactivation of a pathogenicity-associated transposable element triggers major chromosomal rearrangements in a fungal wheat pathogen"

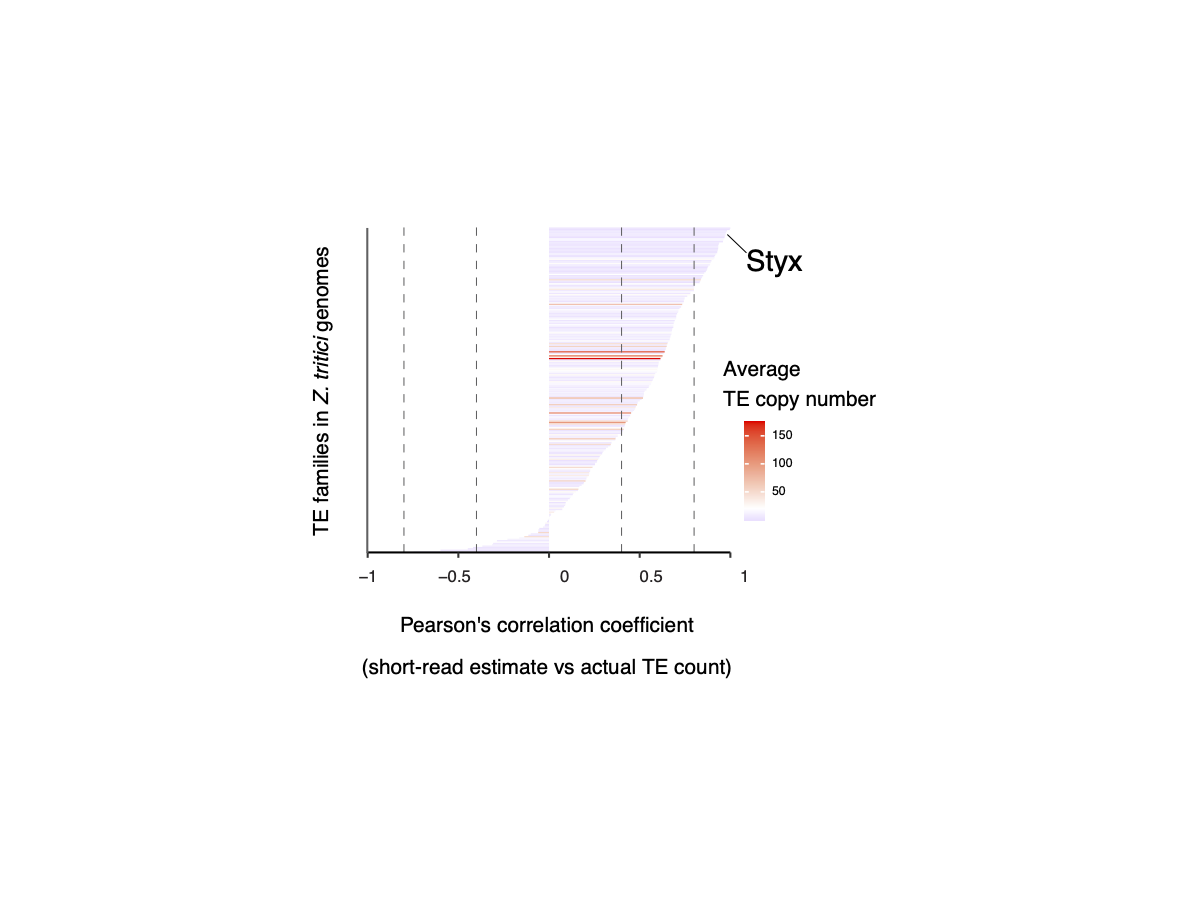

**Figure S1**: Correlation between the estimated copy number from short reads and long reads assemblies of ten *Zymoseptoria tritici* strains. Bars represent the correlation for each of the individual transposable element families identified in the analyzed genomes. The *Styx* element is highlighted and shows among the highest correlations between actual TE count and short-read estimate.

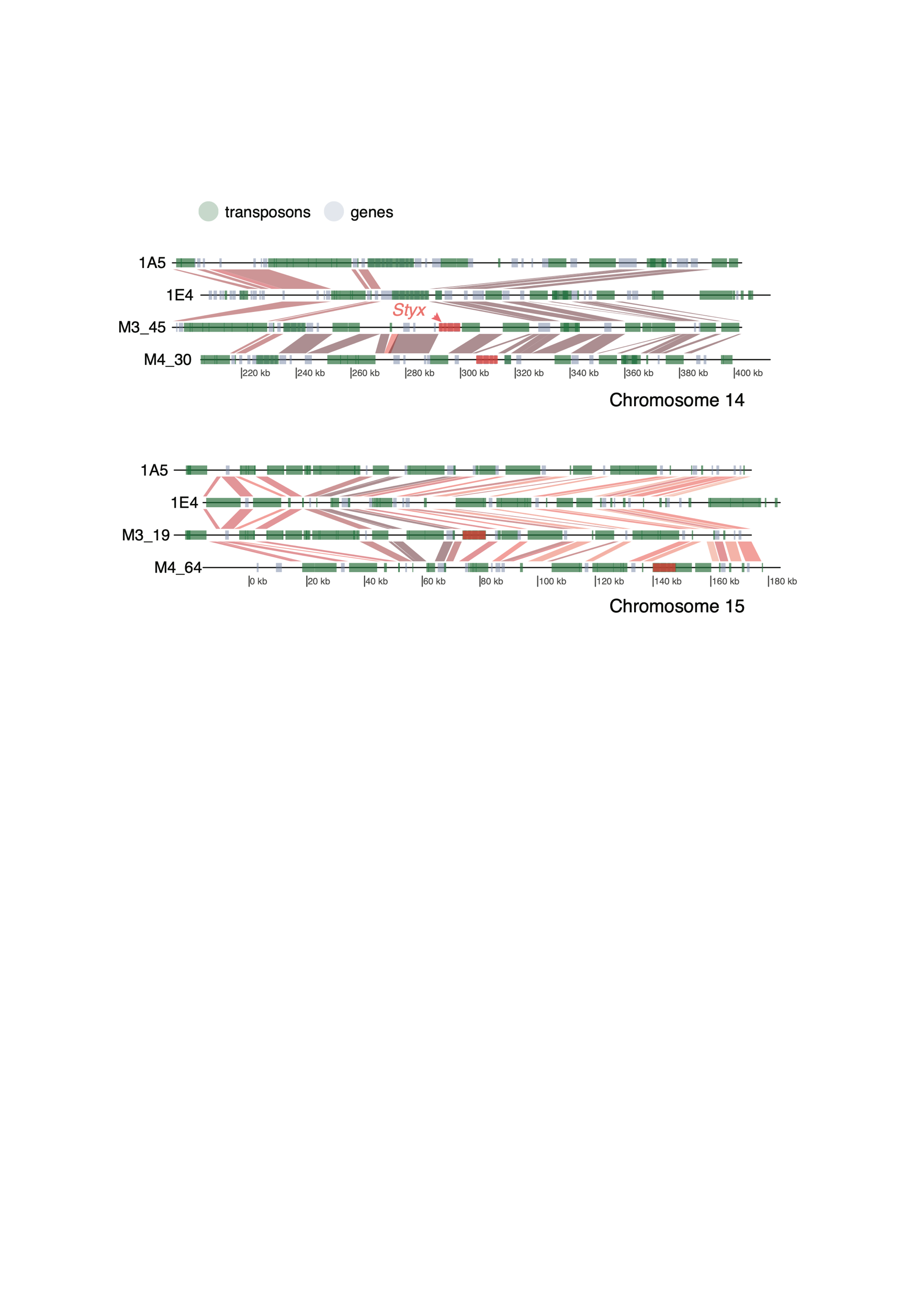

**Figure S2**: Synteny plots illustrating recurrent *de novo* *Styx* insertions in close proximity in the *Zymoseptoria tritici* pedigree. Isolates 1A5 and 1E4 are the parents and M3_45, M3_19, M4_30 and M4_64 are the progeny from the third (M3) and fourth (M4) round of meiosis. *Styx* elements are highlighted in red.

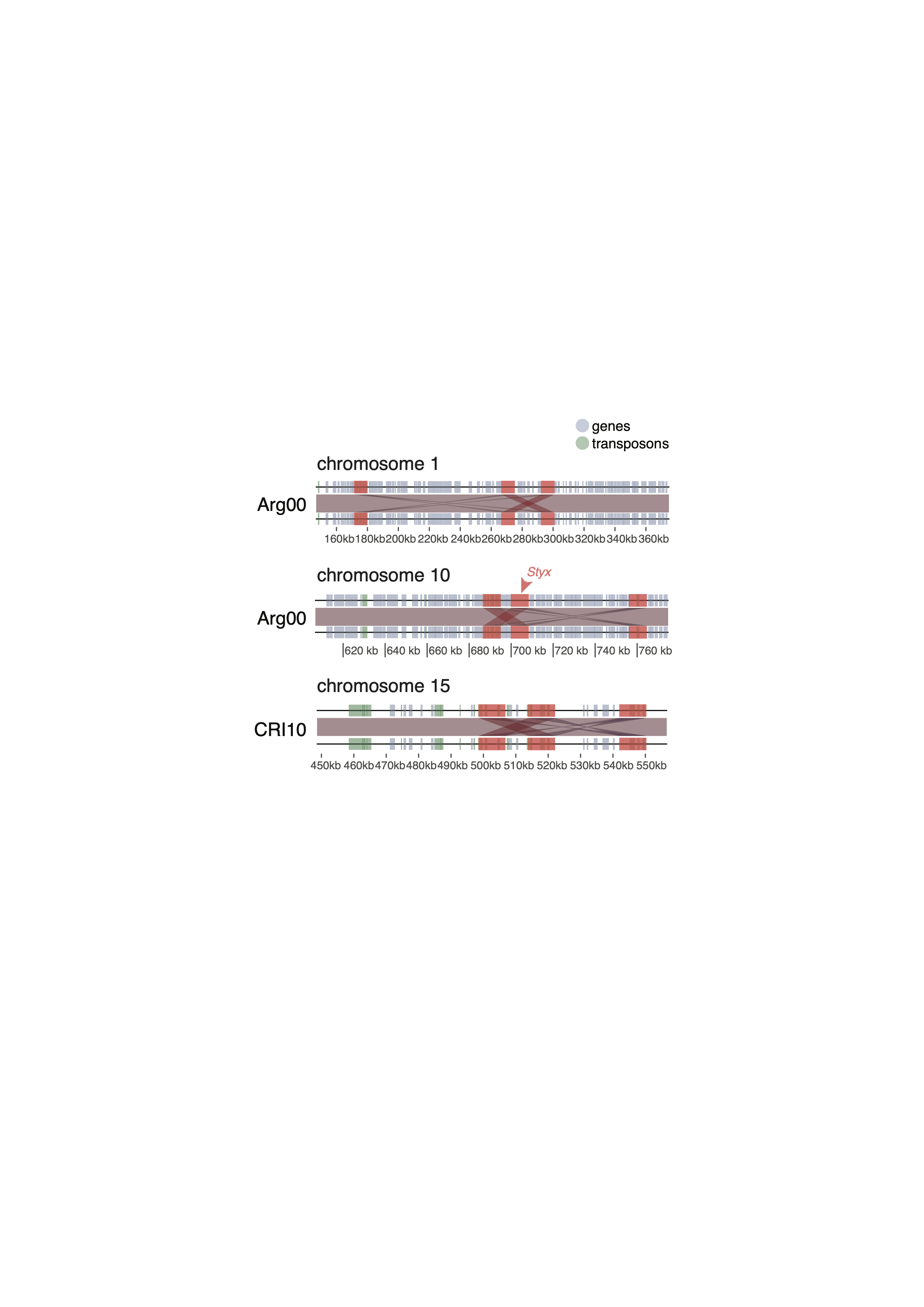

**Figure S3**: Illustration of the recurrent pattern of *Styx* tandem insertions along a same chromosome. Plots show the synteny for the same chromosome and isolate. *Styx* elements are highlighted in red.

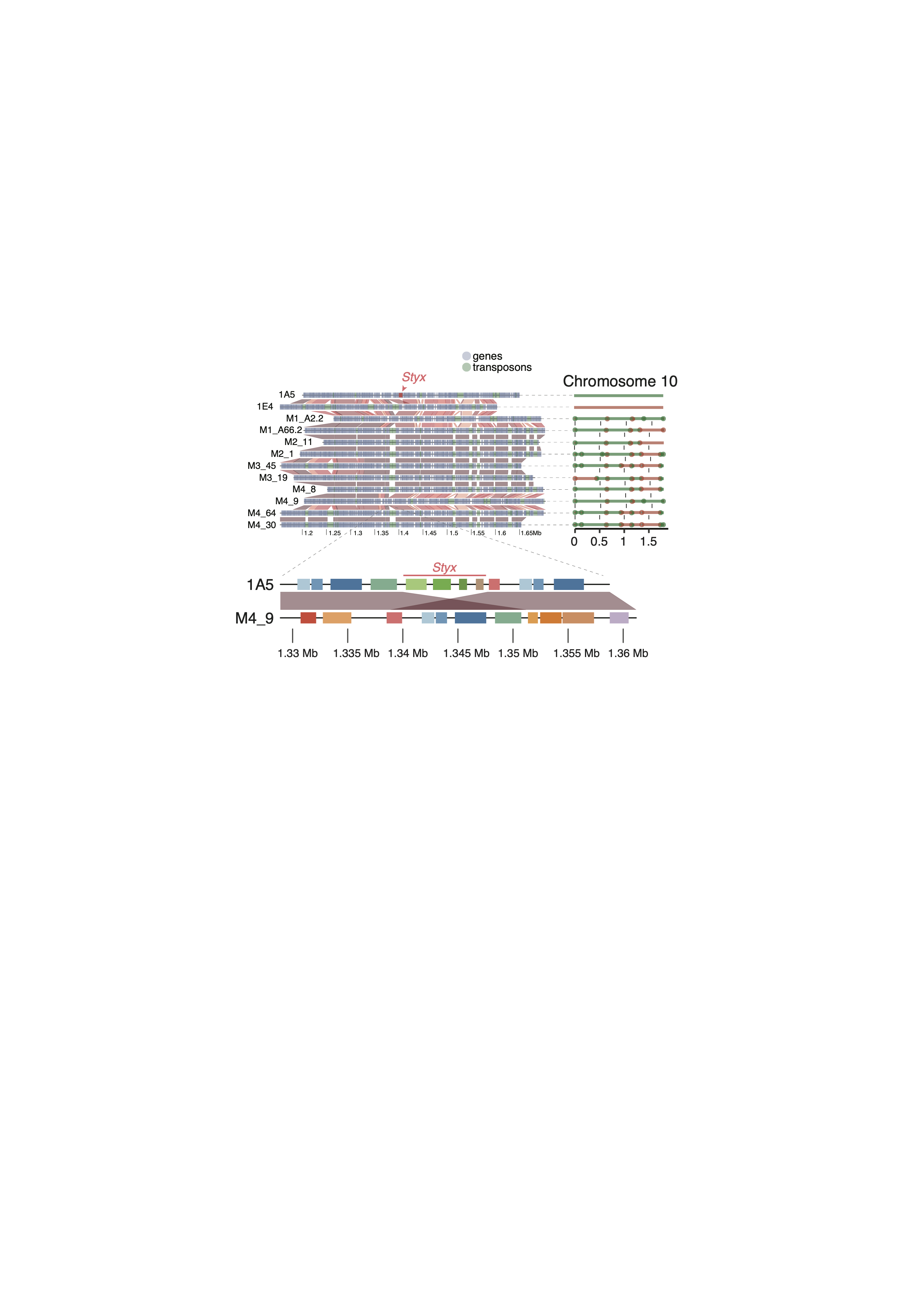

**Figure S4**: Independent excision events of a *Styx* copy on chromosome 10 in the pedigree. Plots show the synteny for the end of chromosome 10 in all progeny isolates. The *Styx* copy present in the 1A5 parent is highlighted in red. Dot-lines on the right side illustrate the recombination events that took place during the crosses. Colours indicate the parental origin of the chromosome segment. The lower synteny plot is a zoom in the excision event in progeny isolate M4_9. The four *Styx* coding sequences are highlighted, and the three surrounding duplicated genes are shown in same shades of blue.
